# Mendelian randomization indicates that TNF is not causally associated with Alzheimer’s disease

**DOI:** 10.1101/673749

**Authors:** Shea J Andrews, Alison Goate

## Abstract

**INTRODUCTION:** Epidemiological research has suggested that inhibition of tumor necrosis factor (TNF)-α in patients with rheumatoid arthritis (RA) reduces the overall risk of Alzheimer’s disease (AD). TNF-α antagonists have been suggested as a potential treatment for AD.

**METHODS:** We used a two-sample Mendelian randomization design to examine the causal relationship between blood *TNF* expression, serum TNF-α levels, and RA on AD risk.

**RESULTS:** Our results do not support a causal relationship between *TNF* expression, serum TNF-α levels or RA on AD risk.

**DISCUSSION:** These results suggest that TNF-α antagonists are unlikely to reduce the risk of AD.

## 1. Introduction

Alzheimer’s disease (AD) is a debilitating neurological condition that is characterized by a progressive deterioration in cognitive function and concomitant functional decline. While the classical neuropathological hallmarks of AD are the deposition of neuritic plaques and neurofibrillary tangles, increasing evidence has highlighted the role of neuroinflammation and microglial innate immune response in AD pathogenesis [1].

Chronic systemic inflammation may also be associated with an increased risk of developing dementia [2], with higher levels of serum proinflammatory cytokines being reported in patients with AD [3]. Chronic systemic inflammation is characterized by the production of the proinflammatory cytokine tumor necrosis factor α (TNF-α) from macrophages. TNF-α is involved in the pathogenesis of chronic autoimmune disorders such as rheumatoid arthritis (RA), but also plays a role in activation of the central innate immune response, including in microglial cells [4]. Inflammation represents a potential means of modifying AD pathogenesis, with the link between peripheral inflammation, TNF-α and neuroinflammation suggesting that TNF-α inhibition may reduce the risk of AD.

AD was found to be more prevalent in RA patients in a nested case-control study of 8.5 million commercially insured adults. Additionally, it was observed that the risk of AD was lower among RA patients who had been exposed to the TNF-α inhibitor etanercept, suggesting that anti-TNF therapy with etanercept could be a potential treatment for AD [5]. The potential role of etanercept as a treatment for AD has gained further media exposure from a recent Washington Post report that analysis conducted by Pfizer, which holds the patent for etanercept outside of the USA, observed a similar decrease in risk of AD in RA patients exposed to etanercept based on insurance claim data [6]. Pfizer, however, elected not to proceed with a clinical trial which was estimated to cost $80 million, according to critics, because etanercept was reaching the end of its patent life, though Pfizer denies this was a factor [6].

While randomized control trials (RCTs) can be reliably used for estimating causal effects, they are expensive to conduct and not all drug targets can be tested in an RCT framework due to ethical considerations or the time-scale involved may be prohibitive. A novel method for estimating the causal effect of drug targets on a disease outcome is Mendelian randomization (MR). MR is a method that estimates the causal effect of an exposure on an outcome by using genetic variants as a proxy for the exposure administered as an intervention in an RCT [7]. MR is analogous to an RCT due to the random allocation of genotypes from parents to offspring at conception (randomization in an RCT) and is thus not affected by reverse causation or confounding variables. In the context of drug development, genetic variants are selected that mimic the action of a drug target, with one allele associated with an altered gene or protein expression (the drug in an RCT) to that of a neutral allele that serves as a reference (the placebo in an RCT) [7]. If the altered genetic allele is associated with a pathway that is causative of the disease, the MR study will detect a change in the clinical outcome. A drug target that has a causative effect on disease is a potential target for drug development, whereas the reverse is true if it is not causative.

In this study, we use Mendelian randomization to evaluate if RA, *TNF* gene expression and TNF-α levels are causally related to AD risk.

## 2. Methods

### 2.1 Instrument Selection

We obtained cis-eQTL data derived from whole blood for *TNF* expression from the eQTLGen project (n = 31,684) [8], pQTL data derived from whole blood TNF-α levels (n = 8,293) [9], and genome-wide significant SNPs for RA from a previous GWAS meta-analysis (14,361 RA cases and 43,923 controls) [10]. To obtain independent SNPs, linkage disequilibrium clumping was performed by excluding SNPs that had an r2 > 0.001 with another variant with a smaller p-value association within 1000kb use a reference panel of European individuals from 1000 Genomes Project (phase 3). Three independent eQTLs for *TNF*, six nominally significant (p < 5e-6) independent TNF-α pQTLs, 56 independent SNPs for RA were selected for analysis. As the effect sizes of the eQTLs were not available in the summary data, the effect sizes were estimated from z-statistics as previously described [11].

The GWAS summary data for AD were from the most recent meta-analysis conducted by the International Genomics of Alzheimer’s Project comprised of 21,982 cases and 41,944 cognitively normal controls (Stage 1 discovery) [12]. The SNPs corresponding to the *TNF* eQTLs, TNF-α pQTLs, and RA SNPs were extracted from the AD GWAS and were harmonized.

### 2.2 Mendelian Randomization Analysis

We used two-sample MR to estimate the causal effect of *TNF* expression, TNF-α levels, and RA on AD. For each variant, we calculated an instrumental variable ratio estimate by dividing the SNP-exposure by SNP-outcome and coefficients. An overall estimate of the causal effect was calculated by combining the individual SNP estimates in a fixed-effects meta-analysis using an inverse-variance weighted (IVW) approach [13]. In order to account for potential violations of the assumptions underlying the IVW analysis, we conducted a sensitivity analysis using MR-Egger regression, which allows all variants to be subject to direct effects [13] and the Weighted Median Estimator (WME), which takes the median effect of all available variants, allowing 50% of variants to exhibit horizontal pleiotropy [13]. Heterogeneity was tested using Cochran’s *Q* statistic [13].

The proportion of variance in the exposure explained by each instrument were calculated as previously described [14]. Power calculations were conducted using the mRnd power calculation tool [15]. All statistical analyses were conducted using R version 3.5.2, with Mendelian randomization analysis was performed using the ‘TwoSampleMR’ package [13].

## Results

The selected instruments for *TNF* expression, TNF-α levels, and RA risk explained 5.93% (F = 285), 1.68% (F = 23.6), and 19.2% (F = 247) of the variance respectively. Given a sample size of 63,926 with the proportion of cases equal to 0.34, this study was adequately powered to detect an OR of any AD of 1.1 for *TNF* expression, 1.19 for TNF-α levels and 1.055 for RA.

There was no evidence of a causal association of *TNF* expression, TNF-α levels or RA on AD risk in the IVW, WME, or MR-Egger regression analyses (Table 2). Similarly, there was no causal association for the individual *TNF* eQTLs. There was evidence of heterogeneity (*Q* = 84.8, *df* = 54, *p* = 0.00472) in RA analysis, but not for the TNF (Q = 3.46, *df* = 2, *p* = 0.177) or TNF-α (*Q* = 2.78, *df* = 5, *p* = 0.733) analysis.

## Discussion

This study examined the causal association of blood *TNF* expression, serum TNF-α levels and RA with AD risk using Mendelian randomization. Despite adequate statistical power to detect an effect, we do not find any evidence that increased *TNF* expression, TNF-α levels or RA risk are causally associated with increased AD risk. These results suggest that TNF-α antagonists, such as etanercept, are unlikely to reduce the risk of AD.

Incidence of AD was reported to be lower in RA patients in a meta-analysis of 10 studies, however, an MR analysis conducted using an earlier AD GWAS also found no causal effect of RA on AD [16]. While animal studies of AD models suggest that TNF-α inhibition ameliorates AD-related pathology, only a few human studies have been conducted [17]. An open-label clinical trial conducted in mild-severe AD patients (n = 15) found that perispinal extrathecal administration of etanercept was associated with significant improvement in cognitive function [18]. In contrast, a double-blind study of etanercept conducted in patients with mild-moderate AD (n = 41) over a 24-week period, found that subcutaneous administration of etanercept showed no effect on cognitive, functional or behavioral assessments [19].

The results of this study should be interpreted in conjunction with its limitations. First, the analysis conducted here was restricted to the expression of *TNF* mRNA in whole blood, the tissue in which the largest eQTL studies have been conducted to date. Analysis in additional tissues may implicate *TNF* expression as a causal risk factor, however, the sample sizes available for other tissues are 30x smaller than that of whole blood and thus have considerably reduced power [20]. Second, the TNF-α GWAS did not contain any genome-wide significant hits, as such, we used nominally significant hits which can result in the inclusion of weak instruments and bias results towards the null. Finally, these MR estimates represent the effect of lifelong exposure to increased *TNF* expression or TNF-α levels, while drugs generally have shorter periods of exposure, and may not distinguish between critical periods of exposure [7].

In conclusion, this Mendelian randomization analysis does not support a causal effect of increased blood *TNF* expression, serum TNF-α levels or RA risk on the risk of AD. These results suggest that, in contrast to recent reports, TNF-α antagonists are unlikely to result in decreased risk of AD. Furthermore, this study highlights how incorporating genetic data into the drug discovery process using Mendelian randomization can improve the drug discovery process.

**Table 1:** Causal effect of *TNF* expression, TNF-α levels and rheumatoid arthritis risk on AD

| | $\beta$ (se) | OR (95% CI) | p |
| --- | --- | --- | --- |
| **TNF - LOAD** |  |  |  |
| rs1121800 | 0.06 (0.05) | 1.07 (0.97, 1.17) | 0.20 |
| rs72855945 | 0.17 (0.19) | 1.19 (0.83, 1.72) | 0.35 |
| rs9469017 | -0.24 (0.17) | 0.79 (0.57, 1.1) | 0.16 |
| IVW | 0.05 (0.05) | 1.05 (0.96, 1.15) | 0.30 |
| WME | 0.06 (0.05) | 1.06 (0.97, 1.17) | 0.21 |
| MR Egger | 0.1 (0.26) | 1.11 (0.67, 1.83) | 0.76 |
| **TNF-α - LOAD** |  |  |  |
| IVW | -0.03 (0.04) | 0.97 (0.89, 1.06) | 0.52 |
| WME | -0.03 (0.06) | 0.97 (0.87, 1.08) | 0.58 |
| MR Egger | -0.07 (0.07) | 0.93 (0.81, 1.06) | 0.34 |
| **RA - LOAD** |  |  |  |
| IVW | -0.01 (0.01) | 0.99 (0.97, 1.01) | 0.37 |
| WME | 0.01 (0.02) | 1.01 (0.97, 1.06) | 0.48 |
| MR Egger | -0.02 (0.03) | 0.98 (0.93, 1.03) | 0.51 |

*TNF*: Tumor necrosis factor mRNA expression; TNF-α : Serum tumor necrosis factor- $\alpha$ levels; RA: rheumatoid arthritis

## Supporting information

Harmonized Instruments

## Funding

SJA and AMG were supported by the JPB Foundation (http://www.jpbfoundation.org).

## Conflicts of Interest

SJA has no conflicts of interest to declare.

AMG served on the scientific advisory board for Denali Therapeutics from 2015-2018. She has also served as a consultant for Biogen, Cognition Therapeutics, AbbVie, Pfizer, GSK, Eisai and Illumina.

